# African origin haplotype protective for Alzheimer’s disease in *APOE*ε4 carriers: exploring potential mechanisms

**DOI:** 10.1101/2024.10.24.619909

**Authors:** Luciana Bertholim-Nasciben, Karen Nuytemans, Derek Van Booven, Farid Rajabli, Sofia Moura, Aura M. Ramirez, Derek M. Dykxhoorn, Liyong Wang, William K. Scott, David A. Davis, Regina T. Vontell, Katalina F. McInerney, Michael L. Cuccaro, Goldie S. Byrd, Jonathan L. Haines, Marla Gearing, Larry D. Adams, Margaret A. Pericak-Vance, ADSP, Juan I. Young, Anthony J. Griswold, Jeffery M. Vance

## Abstract

*APOE*ε4 is the strongest genetic risk factor for Alzheimer’s disease (AD) with approximately 50% of AD patients carrying at least one *APOE*ε4 allele. Our group identified a protective interaction between *APOE*ε4 with the African-specific A allele of rs10423769, which reduces the AD risk effect of *APOE*ε4 homozygotes by approximately 75%. The protective variant lies 2Mb from *APOE* in a region of segmental duplications (SD) of chromosome 19 containing a cluster of pregnancy specific beta-1 glycoprotein genes (*PSGs*) and a long non-coding RNA. Using both short and long read sequencing, we demonstrate that rs10423769_A allele lies within a unique single haplotype inside this region of segmental duplication. We identified the protective haplotype in all African ancestry populations studied, including both West and East Africans, suggesting the variant has an old origin. Long-read sequencing identified both structural and DNA methylation differences between the protective rs10423769_A allele and non-protective haplotypes. An expanded variable number tandem repeat (VNTR) containing multiple MEF2 family transcription factor binding motifs was found associated with the protective haplotype (p-value = 2.9e-10). These findings provide novel insights into the mechanisms of this African-origin protective variant for AD in *APOE*ε4 carriers and supports the importance of including all ancestries in AD research.

## Introduction

Individuals with local African (AFR) ancestral genetic background surrounding *APOE*ε4 have decreased AD risk associated with the APOEε4 allele compared to individuals of European ancestry, thereby demonstrating some natural protective effect in the AFR background. ^1,2^ We recently identified a strong protective interaction between *APOE*ε4 and the A allele of rs10423769, which reduces the AD risk effect of *APOE*ε4 homozygotes by approximately 75% in African Americans (AA), AFR individuals from Ibadan (Nigeria), and genetically admixed Puerto Rican individuals. ^3^ The frequency of the minor rs10423769 A allele is ∼12% in AFR but only ∼0.03% in EUR populations. This protective locus is located 2Mb upstream of *APOE* and lies within a highly segmentally duplicated region of chr19 containing a cluster of pregnancy-specific β-1 glycoprotein (*PSG)* genes as well a long non-coding RNA (ENSG00000282943) (Figure 1). Interestingly, the *PSG* genes are primarily expressed in the placenta with little, if any, expression in the brain. ^4,5^

**Figure 1.**
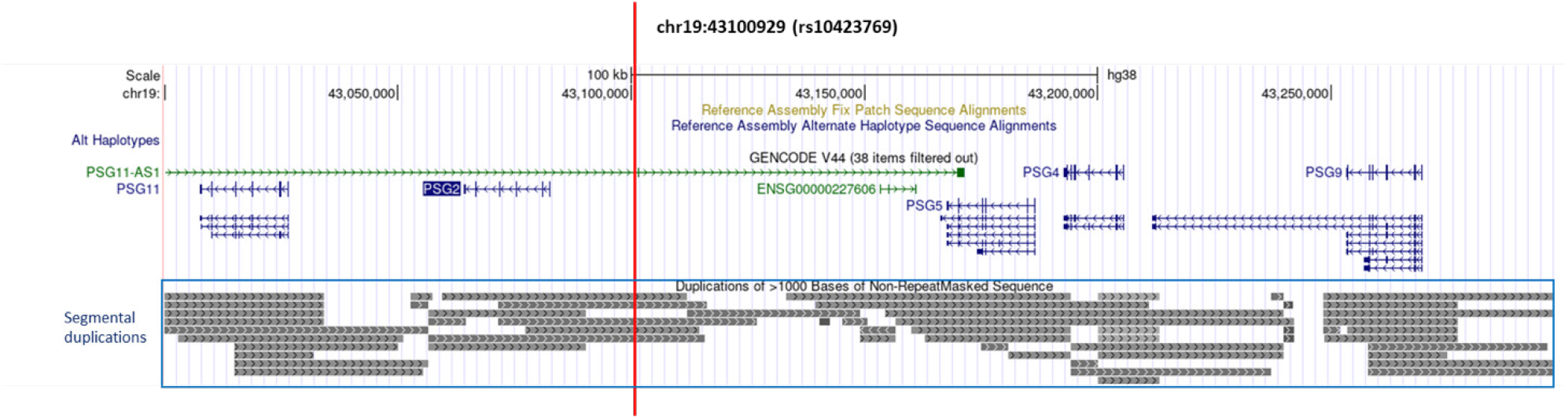
UCSC browser view of surrounding region of rs10423769_A alelle (marked by vertical red line). The annotation of segmental duplication includes all non-allelic intrachromosomal and interchromosomal alignments greater than 1 kb and with more than 90% of sequence identity, excluding common repeats or satellite sequences.

The mechanisms involved in the protective interaction of this locus with *APOE*ε4 are unknown. However, molecular investigation of the protective locus is complicated by its genomic context. The *PSG* region is rich in segmental duplications (SDs) that are difficult to resolve using short-read sequencing data, as traditional sequencing reads are not long enough to be precisely aligned to a specific locus in these repetitive regions. ^6^ In addition, SDs constitute hotspots of recurrent rearrangement by nonallelic homologous recombination, resulting in high occurrence of copy number variations (CNVs), gene conversion, and structural variants (SVs) ^7,8^. SVs could affect chromatin structure, which we have previously described as differing between AFR and EUR brain astrocytes. ^9,10^ In addition, the SD region of the *PSG*s has been previously reported to have many deletions and duplications that vary between ancestries. ^4^

To gain insight into its mechanisms of protection, we performed an initial genetic characterization of this protective locus. First, we refined the haplotype harboring the protective locus using a large, short read-sequencing database, including individuals from diverse ancestries from the Alzheimer’s Disease Sequencing Project (ADSP). Second, we used long-read sequencing to confirm the location of the haplotype in this highly segmentally duplicated region of chromosome 19. Third, we used the long-read sequencing data to identify the SVs and DNA methylation profiles in the *PSG* region. Finally, we explored an expanded variable number tandem repeat (VNTR) enriched in multiple MEF2 family transcription factors binding motifs present in this genomic region. These findings provide novel insights into the potential mechanisms underlying this AFR-origin protective variant for AD in *APOE*ε4 carriers. Although biologically complex, a better understanding of the genetic and molecular factors involved in the protection against risk of *APOE*ε4 driven by this region presents a therapeutic opportunity for all ancestries.

## Methods

### Haplotype block analysis

We extracted phased genotype data from rs10423769_A allele carriers from the Alzheimer’s Disease Sequencing Project Release 4 (ADSP R4: ng00067.v10) of short-read whole genome sequencing (Web resources). In the region of 70 kb surrounding rs10423769, we selected all SNPs with a MAF ≥ 0.05 and Hardy–Weinberg equilibrium (HWE) exact test p-value > 0.001 in the African population of the 1000 Genomes project (1000G) dataset for analysis (n=301). ^11^ Haploview 4.2 software was used to define the haplotype blocks. ^12^ Plink v1.90 ^13^ (Web resources) was used to calculate Linkage Disequilibrium (LD) using ADSP R4. LDhap from the LDlink (5.6.5 Release) web-based application ^14^ (Web resources) was used to map and calculate frequencies of the haplotypes harboring the rs10423769_A allele across population groups from the 1000G Phase 3.

### Long read whole genome sequencing (LRWGS)

LRWGS was generated from DNA extracted from either cerebellum samples excised from frozen brain or peripheral blood samples from 38 individuals that were heterozygous or homozygous for rs10423769_A, or non-carriers, as described in Supplementary Table 1. Brain samples were obtained from the biorepository of the John P. Hussman Institute for Human Genomics (HIHG) and Brain Endowment Bank at the University of Miami, as well as Emory University Goizueta Alzheimer’s Disease Research Center (GADRC) Brain Bank. Blood samples were obtained from participants as part of ongoing research projects in studying Alzheimer’s disease in individuals of African ancestries (AG052410 and AG072547, P.I. M. Pericak-Vance).

DNA was extracted in the HIHG biorepository using the AutoGen FlexSTAR using standard procedures without further size selection. Libraries were constructed using the SQK-LSK109 ligation kit from Oxford Nanopore Technology (ONT). Samples were loaded onto PromethION R9.4.1 flow cells and sequenced in 72-hour data acquisition runs on the PromethION24 device. Base calling was performed with Guppy version 3.3.2 which simultaneously produces MM and ML methylation tags in the unaligned bam file. Resulting bam files were then converted to fastq files (samtools v1.2) preserving these tags in the meta data for each read). Resulting FASTQs were aligned to GRCh38 using minimap2 v2.17-r941 where the methylation tags were preserved from the previous step. Small variant calling was performed with Clair3 (v1.0.3). Sniffles2 (v2.0.7) was used for structural variant calling (default parameters were used individually on each sample, then a joint call was performed with default parameters with sniffles2). Aligned BAM files were examined using the Integrative Genome Viewer v.2.4.10 (Web resources) in Third Gen quick consensus mode.

### Local assembly and motif finding

TREAT (Tandem REpeat Annotation Toolkit) assembly tool with Otter ^15^ (Web resources) was used for local assembly of tandem repeat regions in chr19 and annotation of VNTRs. The association between rs10423769 allele and the tandem repeat region length measured in bp was evaluated using Wilcoxon rank sum test. MEME Suite ^16^ (Web resources) was used for de novo motif finding and the algorithm FIMO (version 5.5.5) ^17^ within MEME Suite was used to identify known transcription factors (TF) binding motifs in the region. We screened defined transcription factors binding site databases JASPAR CORE 2022 (vertebrates non-redundant) ^18^ and selected those binding sites with q-value < 0.001.

### Allele-specific differential methylation analysis

BAM files with methylation tags were phased by Longshot v0.4.5 ^19^ using a region of +/-40 kb from rs10423769 with a minimum coverage of 10 and minimum alternative allele fraction of 0.35. Allele-specific bedmethyl files with aggregation of modified bases were obtained with the tool modkit v0.1.12 (Web resources) with the options --partition-tag HP, --combine-strands, --cpg, and --ignore h. The DMA module from Nanomethphase v 1.2.0 ^20^ were used for differential methylation analysis with -smf set to FALSE.

## Results

### Haplotype block analysis (based on short-read ADSP data)

We determined the composition of the haplotype harboring the rs10423769_A using the ADSP R4 set of short read whole genome sequencing. We identified 1,962 individuals carrying at least one rs10423769_A allele. Six haplotype blocks were identified in the region surrounding rs10423769_A (Figure 2). No recombination is observed between the most frequent haplotypes (∼53%) in block 1, block 2 (harboring the rs10423769_A) and the remaining blocks 3-6. Supporting this, rs10423769 has a D’ > 0.95 (LOD ≥ 2) with all SNPs from these six blocks, with the exception of two positions (rs8107144 and rs7250796) with D’ < 0.3. Thus, we determined that the minimum shared haplotype harboring rs10423769_A allele was approximately 21kb (chr19:43099521-43120243) spanning six haplotype blocks. An extended haplotype including blocks 7-9 (chr19:43121359-43132912) was also identified with a D’ = 0.76 between blocks 6 and 7.

**Figure 2.**
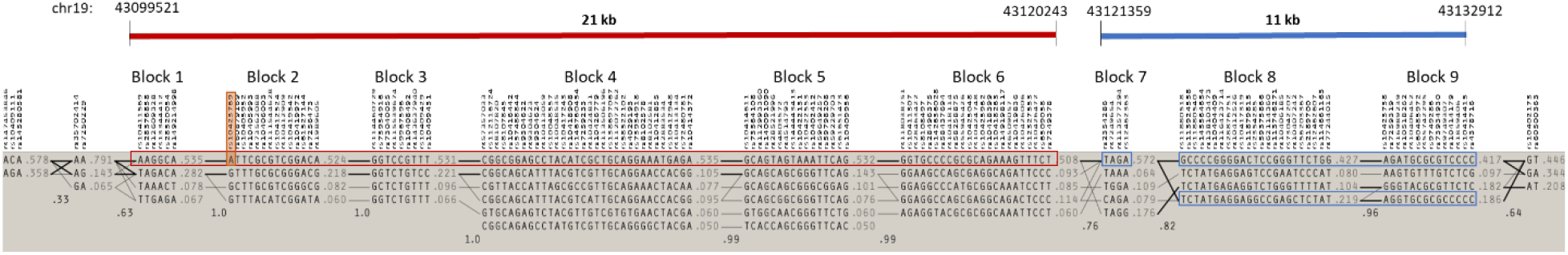
Haplotype blocks identified in 1,962 individuals carrying at least one rs10423769_A allele using Haploview 4.2 and ADSP. Lines between blocks indicate co-occurrence of adjacent haplotypes in individuals with line thickness representing frequency of co-occurrence across individuals. Haplotype block frequencies are shown in the right of each block (≥ 0.05). Multiallelic D’ is shown on the bottom of crossing areas, which represents the level of recombination between blocks. Blocks with D’ > 0.8 were considered the same haplotype. Rs10423769 is marked by an orange box. The 21kb (chr19:43099521-43120243) minimum shared haplotype is marked by a red box and the 11 kb extended haplotype (chr19:43121359-43132912) is marked by blue boxes.

In order to support these findings, we performed LD analysis using r^2^ statistics and the entire ADSP R4 dataset of ∼36,000 individuals, which showed almost perfect linkage (r^2^ >0.95) between rs10423769_A and 13 markers distributed along the six blocks of the 21 kb minimum shared haplotype (Supplementary Table 2). Eleven markers distributed along the 11 kb extended haplotype showed r^2^ >0.75.

### Frequency of the haplotype across populations

Using the allele frequency information from 1000G to verify the population-specific frequency of the 21 kb minimum shared haplotype, we found the same haplotype across multiple admixed populations containing African ancestry (African American, Afro-Caribbean, and Puerto Ricans admixed populations) (Figure 3), but not in individuals of Mexican, Peruvian, East Asian, South Asian or European populations. Interestingly, the haplotype frequency was similar between West (Nigeria) and East (Kenya) populations.

**Figure 3.**
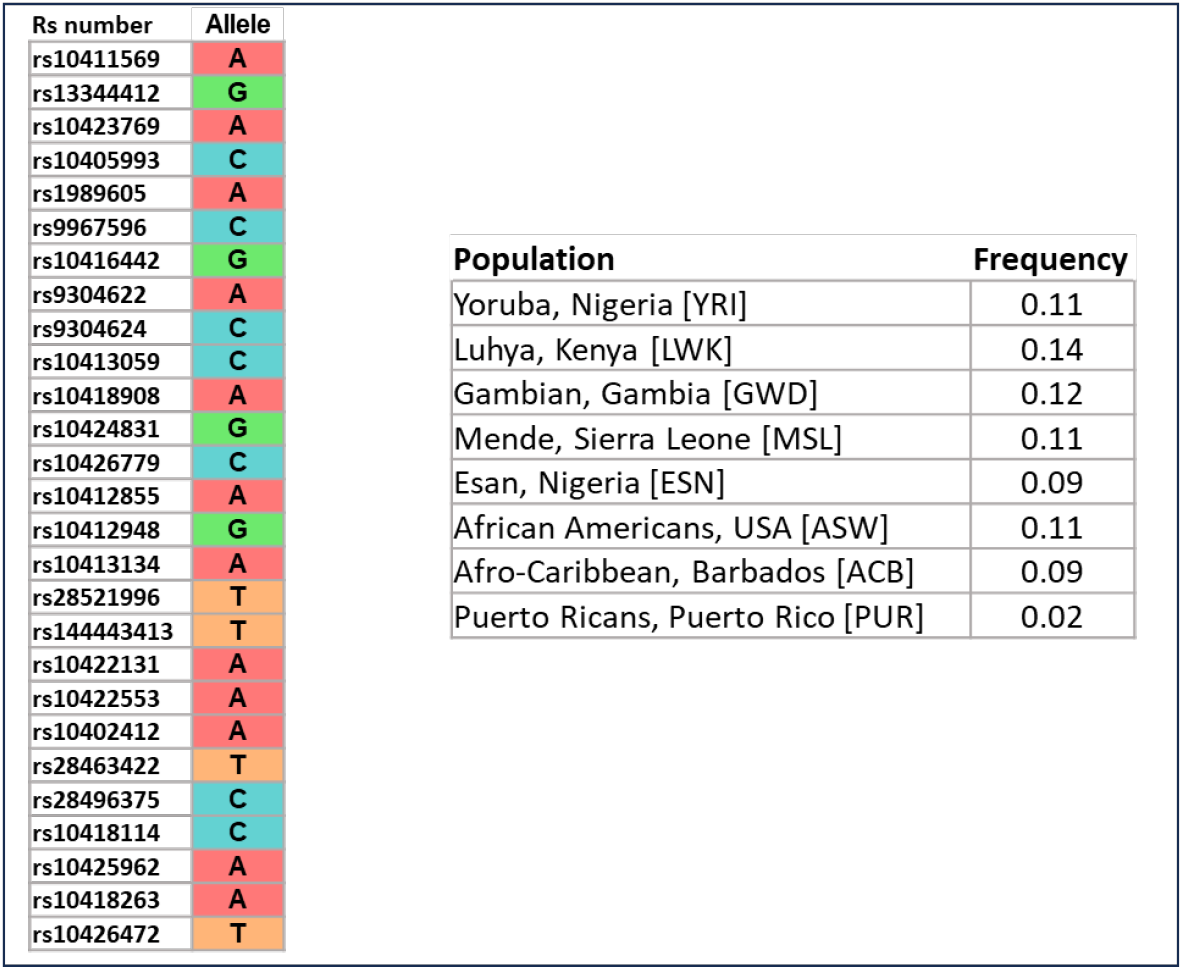
Population-specific frequency of the minimum shared haplotype harboring the rs10423769_A allele in 1000G. LD pruned haplotype across 10 blocks determined by Haploview are shown. Frequencies were calculated with the tool LDhap from LDlink web-based application.

### Validation of minimum shared haplotype with LRWGS

Since rs10423769_A allele is in an area of segmental duplication, LRWGS was used to validate whether rs10423769 is a true variant or an artifact of incorrect mapping in the SD region. LRWGS of 38 brain and blood samples including rs10423769_A homozygous, heterozygous and non-carriers (Supplementary Table 1) was performed with ONT yielding an average genome coverage of 21.5 ± 5.3X and average read length of 10.4 ± 3.0 kb. The average depth of high-quality reads (MAPQ ≥ 60) for the rs10423769_A position (chr19:43100929) was 16.0 ± 4.9X and all reads mapped uniquely for this specific position. In addition, analysis of potential secondary alignments of reads spanning the 21 kb minimum shared haplotype confirmed that the variant pattern identified along the haplotype block are specific to this region and not found contiguously in any other region of the genome (Figure 4). The frequency of variants in almost perfect linkage disequilibrium (r^2^ >0.95) were confirmed in the LRWGS variant calling (Supplementary Table 2).

**Figure 4.**
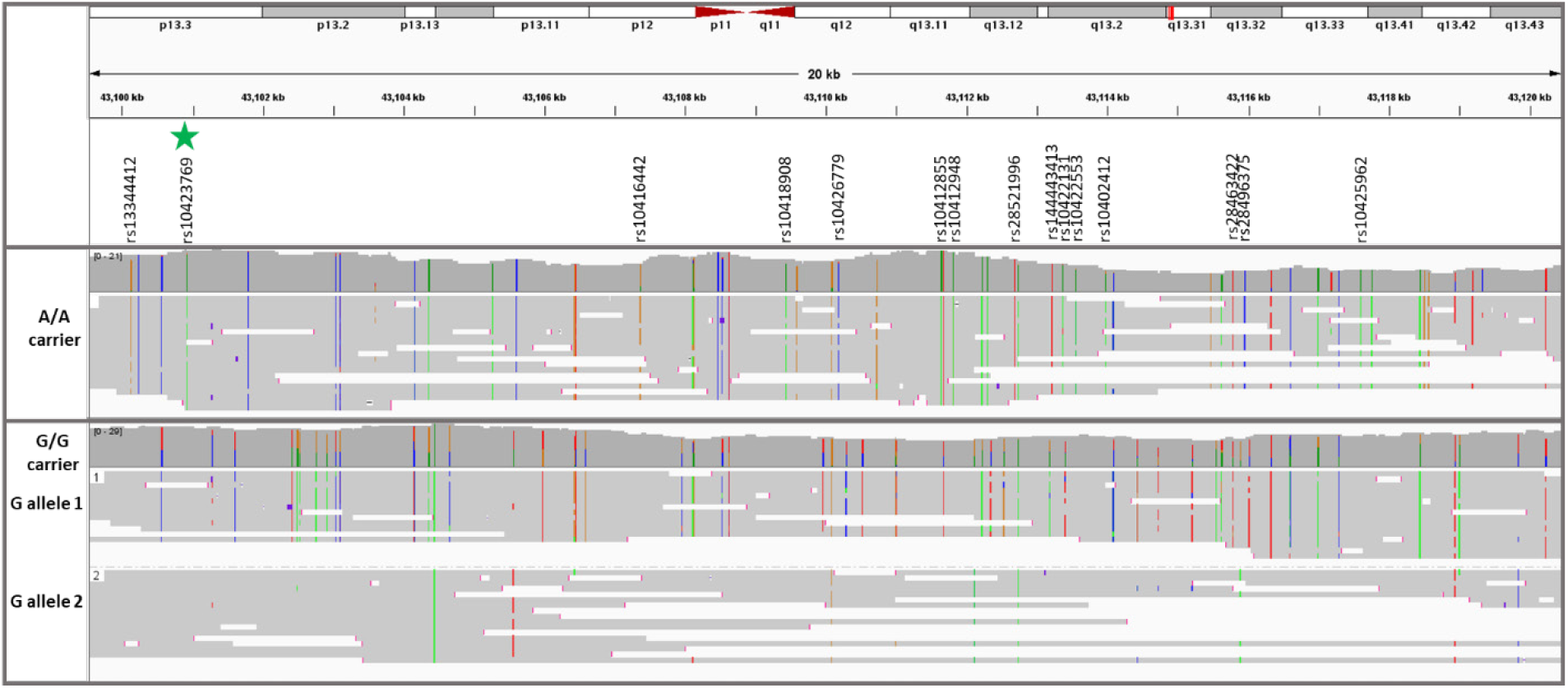
Read the coverage and map of the 21 kb minimum shared haplotype from two representative individuals homozygous for either the rs10423769_A or rs10423769_G alleles. The green star marker represents the rs10423769_A allele, and all other listed markers are in LD>0.95 with rs10423769_A. Colored lines represent IGV consensus SNPs: A, green; C, blue; T, red; G, orange.

### Analysis of structural variation

After mapping the haplotype associated with rs10423769_A, we explored other genomic and epigenomic features that could contribute to the functional mechanisms involved in the protection afforded by this locus. The LRWGS data were used to detect germline SVs in the 2 Mb region (chr19: 43000000-45000000) spanning from the protective locus to *APOE*. Sniffles2 identified 87 SVs in the region, including 42 deletions, 43 insertions, and two breakends. One insertion located at chr19:43132126 (32 kb from rs10423769) was identified with a frequency of 0.80 in rs10423769_A homozygotes and 0.07 in non-carriers. This insertion is located in an annotated repetitive region (772bp, chr19:43,131,850-43,132,621) (Web resources) approximately 32 kb from the protective locus, on block 9 of the expanded haplotype (Figure 2). Further investigation of the 772bp region of the reference genome indicates the presence of a VNTR with repetitions of a 29 bp tandem repeat pattern. Local assembly of this region revealed that the A allele is associated with expanded VNTR alleles (p-value = 2.944e-10, Figure 5) containing a higher number of the 29 bp tandem repeat. We analyzed the region for the presence of motifs and known TF binding sites and identified that the 29 bp repetitive sequence carries predicted binding sites motifs for the MEF2 family of transcription factors. MEF2D had the highest score by FIMO motif finding tool, ^17,18^ followed by MEF2B and MEF2A (Figure 6).

**Figure 5.**
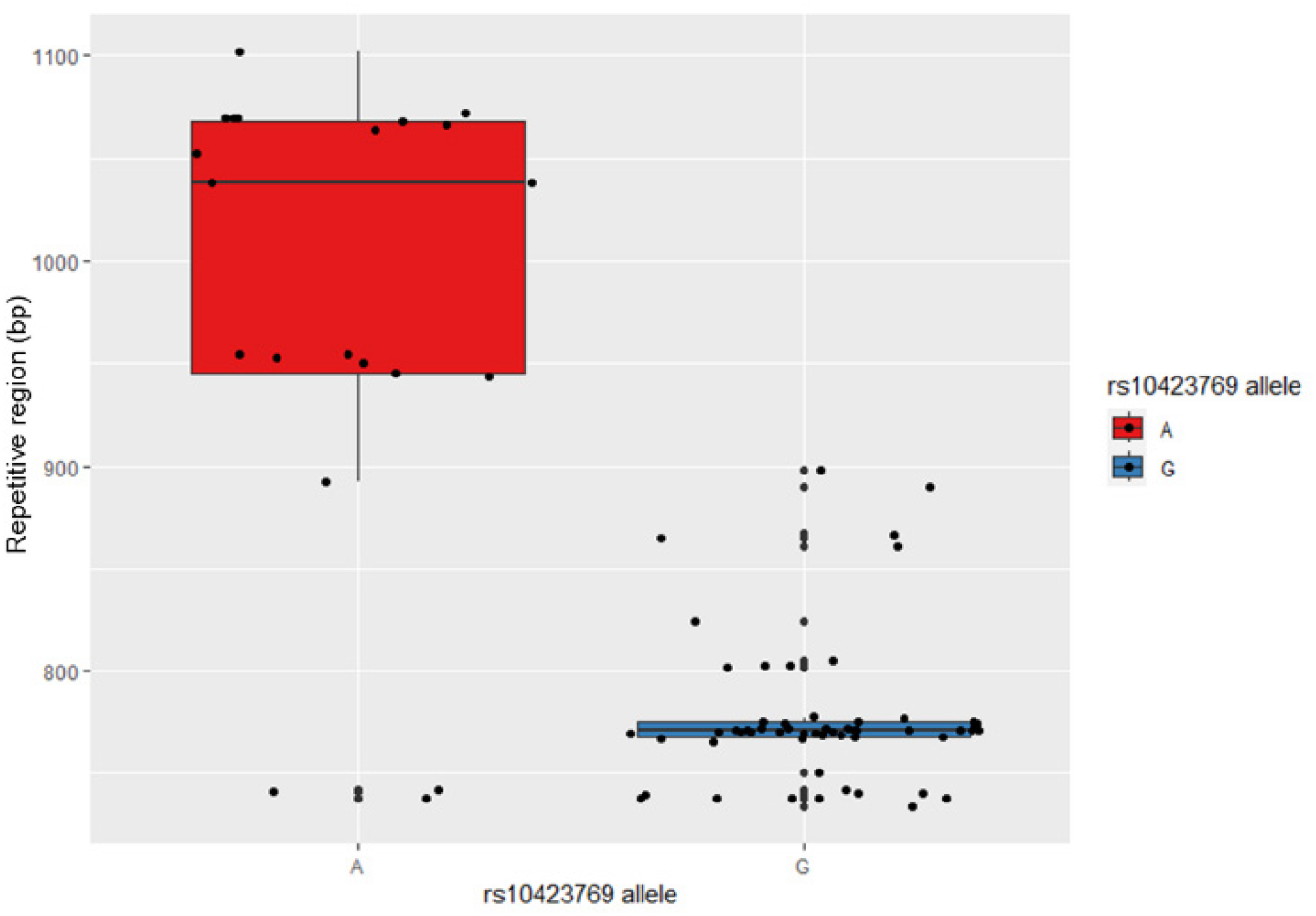
VNTR length correlation with rs10423769 haplotype (p-value = 2.944e-10).

**Figure 6.**
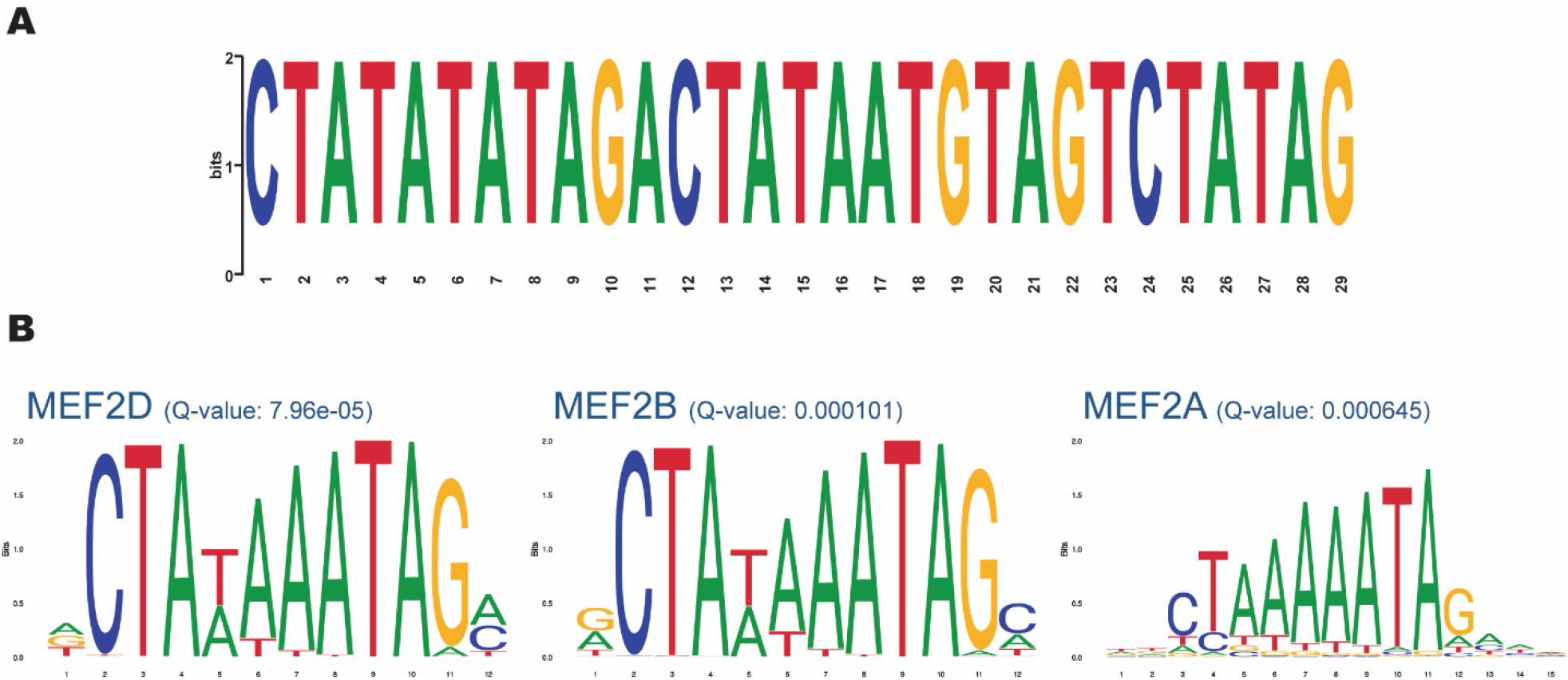
Motif analysis of the expanded VNTR allele associated with rs10423769_A. **A.**29 bp Repetitive sequence identified by MEME Suite ^16^. **B**. MEF2 family of TF binding motifs with FIMO motif finding tool q-value ^17,18^.

### Differential methylation analysis

We also hypothesized that the mechanism of protection could be related to differential DNA methylation at the haplotype. Thus, we performed allele-specific methylation analysis of five heterozygote rs10423769_A/G brain samples to evaluate methylation differences in the 21 kb minimum shared haplotype and surrounding region. We identified 17 differentially methylated positions (DMP) (FDR < 0.01) comparing rs10423769_A to rs10423769_G haplotypes. Further analysis indicated that differences in methylation on those positions occurred due to base changes in the haplotype sequences between rs10423769_A to rs10423769_G haplotypes, with gain or loss of CpG sites (Figure 7).

**Figure 7.**
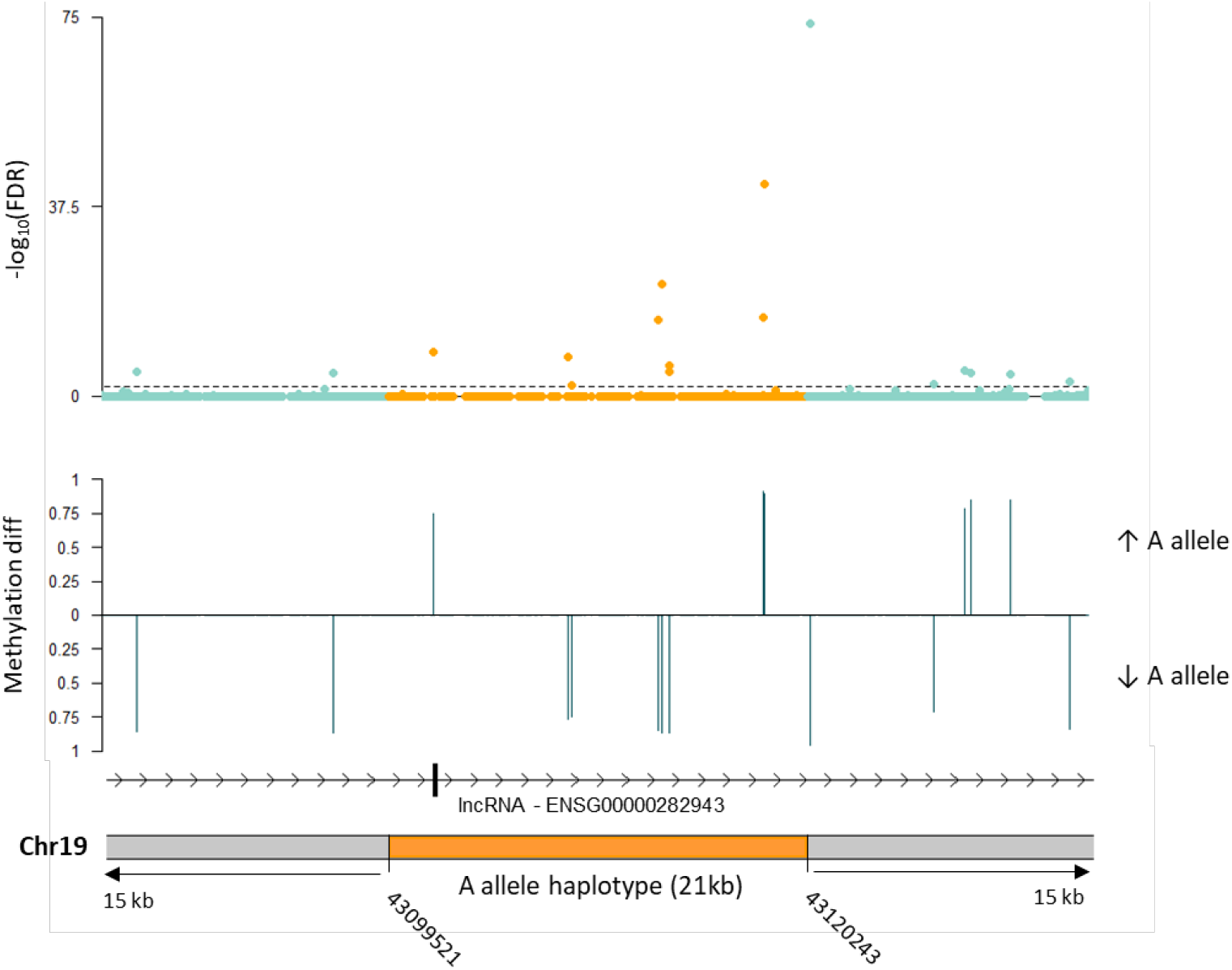
Allele-specific differential methylation analysis in the rs10423769_A allele 21 kb minimum shared haplotype and surrounding region using 5 brain samples heterozygous for rs10423769. The dotted line indicates FDR <0.01. “Methylation diff” refers to the difference in mean methylation levels between the rs10423769_A and rs10423769_G haplotypes. “Methylation diff” is shown for DMP with FDR < 0.01.

## Discussion

Herein we report the initial efforts to characterize the protective locus tagged by the A allele of rs10423769 which reduces the AD risk effect in AFR ancestry *APOE*ε4 homozygotes by approximately 75%. Understanding the mechanisms involved in the protective effect is challenging, since the variant is located 2 Mb away from *APOE*, in a large area of SD containing a *PSG* gene cluster composed of 10 genes (*PSG1*-*9, PSG11*). Given the SD in the region, one of the first questions to answer was whether the locus was unique or duplicated. We demonstrate that it is a unique feature lying in a large area of segmental duplications. We confirmed that the minimum shared haplotype is from AFR origin and is found in all AFR populations represented in 1000 Genomes and admixed AFR populations, but not in Mexican ancestry, Peruvian, East Asian, South Asian and European populations. The presence of the haplotype in both Western and Eastern African populations suggests it is likely old in its AFR origin.

The haplotype of ∼21kb shared by all rs10423769_A carriers and the extended haplotype of another 11 kb in high LD with rs10423769_A (r^2^ = 0.75) overlap with the lncRNA (*PSG11-AS1*/ ENSG00000282943) (Supplementary figure 1). Overall, this lncRNA is expressed in all tissues at very low levels in the cerebral cortex according to The Genotype-Tissue Expression (GTEx) Project Release V8 (Web resources), with cerebellum, cerebellar hemisphere, and cultured fibroblasts having higher levels of expression. ^21^ In contrast, the *PSG* genes belong to family of glycoproteins that are primarily expressed in the human placenta. ^4,5^ Interestingly, data from the Allen Institute for Brain Science (Web resources) suggest that *APOE* has its highest overall expression just before and after birth. More studies will be needed to elucidate any potential involvement of the *PSG* genes or lncRNA directly in the protective effect on *APOE*ε4.

The *PSG* locus was previously reported to have a higher frequency of copy number variations, deletions and duplications, compared to the genome average, with differences in frequency and distinct breakpoints between AFR and non-AFR haplotypes. ^4,22^ However, given the duplication pattern in this region, the reliability of such reports is uncertain. Thus, we used LRWGS around the SD to characterize the region and the protective and non-protective haplotypes. We did not see obvious evidence of the SV patterns previously reported. This may be because of the technological limitations of these previous reports that cannot account for potential changes in segmental duplication patterns between individuals that may be reported as SVs. Gaining a better understanding of the genomic structure of this locus is critical since it is possible that differences in SVs or pattern of SDs between the protective and the risk haplotypes could influence chromatin interactions or other regulatory mechanisms in the area. In fact, it has been shown that chromatin reorganization happening during cellular aging leads to the re-expression of *PSG* genes, ^23^ suggesting a possibility of differences in SV affecting *PSG* gene expression. While the depth of reads allowed us to validate the structure of the protective haplotype, it was not enough to allow assembly of the entire 0.55 Mb SD region. This will require a much higher read depth, perhaps as high as 100x.

Several neurodegenerative diseases have been associated with VNTRs ^24^ in general and AD specifically. For example, increased length of a 25 bp repeat unit located in intron 18 of *ABCA7* was associated with increased AD risk. ^25^ Interestingly, the protective haplotype was associated with expanded VNTR alleles which are enriched for a 29 bp motif with multiple MEF2 binding motifs. These types of clusters and VNTRs can be found in many areas of the genome, but what is compelling here is the significantly larger VNTR associated with the protective haplotype vs. the non-protective haplotype. The MEF2 family of TF have an important role during both development and adulthood, participating in neuronal development, synaptic plasticity, cognitive reserve and neurodegenerative diseases, by controlling the expression of several genes and miRNAs ^26^. The expression of all MEF2 isoforms is high in the brain, with the expression of MEF2A and MEF2D increasing with neuronal differentiation and maturation, whereas the expression of MEF2C remains relatively stable throughout. ^27^ Barker et al. (2022) found that MEF2 transcriptional network demonstrated the strongest association with predictive good cognition towards the end of life. Overexpression of *MEF2A/C* in a mouse model of tauopathy had positive effects on cognitive flexibility. ^28^ However, other authors suggest MEF2 has a negative effect on memory function. ^29^ Thus, depending on the interaction with other co-factors, such as chromatin-modifying enzymes or polymerase complex, and the cell type, MEF2s can either activate or suppress gene expression ^27^. It was reported that a MEF2A variant (p.Pro279Leu), which decreases MEF2A’s function in transcriptional activity, was significantly enriched in LOAD patients. ^30^ In addition, elevated methylation at an enhancer region of *MEF2A* that reduced *MEF2A* expression has been reported in AD, ^31^ further linking MEF2 decreased activity with AD. Further studies are needed to investigate if and how this VNTR may contribute to lowering *APOE*ε4 risk.

Differences in methylation status between AD cases and controls have been noted, including in the *APOE* region. ^32–34^ We identified several DMP in the protective haplotype, 2 Mb away from *APOE*. Long-range effects of methylation as far as 10 Mb from promoter regions have been documented to play a role in regulation of gene expression. ^35^ Therefore, further follow-up of these DMPs is needed to determine the role of these alterations on *APOE* expression specifically or other genes in the region, and on AD risk in general.

One important question to be addressed in the future is whether the protective association of rs10423769_A with *APOE*ε4 involves lowering of *APOE*ε4 expression. Single nuclei RNA-sequencing data from our group suggests that the expression of *APOE* is much lower in one individual who is an *APOE* ε4/ε4 carrier and homozygote for rs10423769_A when compared to rs10423769_G carriers. ^9^ Efforts are underway to identify brain material carrying both the rs10423769_A allele and APOE4, but the low availability of tissue from the African ancestry population makes this more challenging.

Overall, while dozens of risk variants for AD have been described over the last decade, protective variants for AD have received less attention. Through evolution, protective mechanisms, usually with minimal side effects, have been established naturally. Increasing the number of studies on these protective variants to understand their processes is essential for the advancement of therapeutics in AD. In addition, our study illustrates the importance of including diverse populations in genetics studies to ensure broad representation and open opportunities to uncover and understand Alzheimer disease and biological mechanisms from a wider perspective.

## Data and Code Availability

Short read WGS from the ADSP is available via NIAGADS (ADSP R4: ng00067.v10) (Web resources). Long read sequencing is available upon request from the corresponding author. No custom code was created for this manuscript. All data processing was performed using publicly available software as referred to in the Methods.

## Acknowledgments

This research was supported by the National Institute on Aging through the following grant numbers: U01AG072579, RF1AG059018, R01AG070864, R01AG072547, U01AG066767, P30AG066511, and U01AG052410. Additional funding was received by a Zenith Award (Alzheimer’s Association, J.M.V), the BrightFocus foundation (A2018425S, J.M.V) and Alzheimer’s Association Research Fellowship to Promote Diversity (AARFD-24-1309441, L.B.N.). We are grateful to the many participants, researchers, and staff who contributed significantly to this study.

## Web resources

Alzheimer’s Disease Sequencing Project Release 4, https://adsp.niagads.org/

Plink v1.90, www.cog-genomics.org/plink/1.9/

LDhap and LDlink, https://ldlink.nih.gov/

Integrative Genome Viewer (IGV), http://software.broadinstitute.org/software/igv/

TREAT (Tandem REpeat Annotation Toolkit), https://github.com/holstegelab/treat/

MEME Suite, https://meme-suite.org/meme/tools/meme/

FIMO, https://meme-suite.org/meme/tools/fimo/

Modkit, https://github.com/nanoporetech/modkit/

Tandem repeat annotation, https://github.com/PacificBiosciences/pbsv/tree/master/annotations/

BrainSpan Atlas of the Developing Human Brain (Allen Institute for Brain Science), https://www.brainspan.org/rnaseq/search/index.html

Genotype-Tissue Expression (GTEx), https://www.gtexportal.org/

