## Supplementary Tables and Figures for "African origin haplotype protective for Alzheimer’s disease in *APOE*ε4 carriers: exploring potential mechanisms"

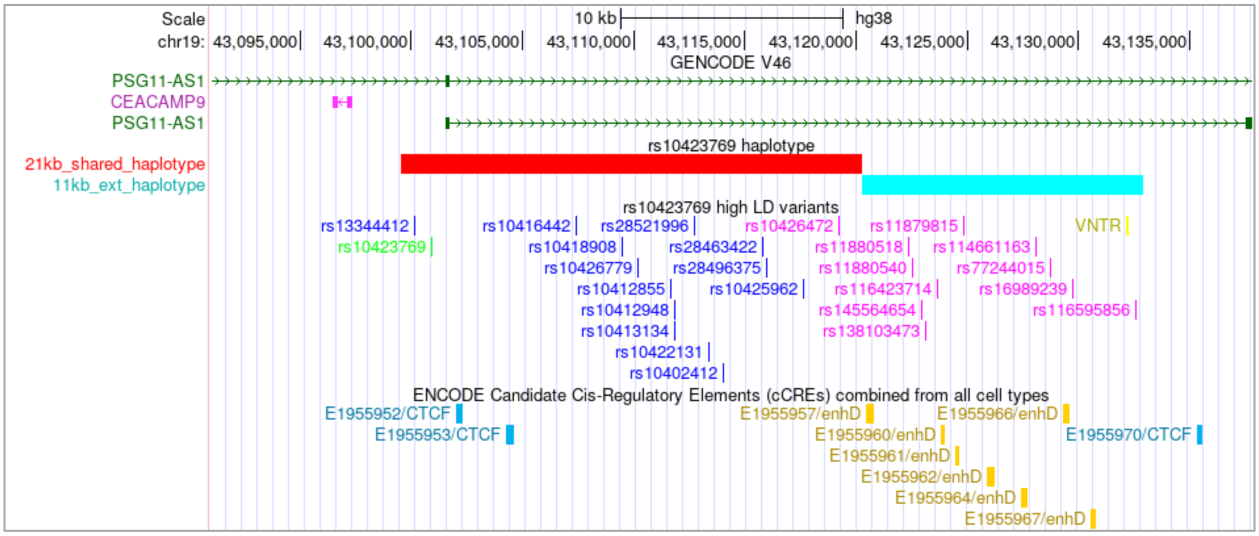


**Supplementary Figure 1.** UCSC browser view (hg38) of 21 kb minimum shared haplotype (in red) and the 11 kb extended haplotype (in light blue) and the ENCODE Registry of candidate cis-Regulatory Elements (release September 14, 2018). The protective allele rs10423769 is shown in green. SNPs with r2 >0.95 with rs10423769_A are shown in blue and SNPs with r2 >0.75 with rs10423769_A are shown in purple. Tandem repeat region (VNTR, chr19:43132126-43132260) is shown in yellow.

**Supplementary Table 1.** Tissue and rs10423769 genotype for 38 samples submitted to LRWGS

| Sample number | Type | Research Center | rs10423769 |
|---|---|---|---|
| sample1 | brain | UM/BEB | A/G |
| sample2 | brain | UM/BEB | A/G |
| sample3 | brain | UM/BEB | A/G |
| sample4 | brain | UM/BEB | A/G |
| sample5 | brain | Emory | A/G |
| sample6 | blood | UM | A/A |
| sample7 | blood | UM | A/A |
| sample8 | blood | UM | A/A |
| sample9 | blood | UM | A/A |
| sample10 | blood | UM | A/G |
| sample11 | blood | UM | A/G |
| sample12 | blood | UM | A/G |
| sample13 | blood | UM | A/G |
| sample14 | blood | UM | A/G |
| sample15 | blood | UM | A/G |
| sample16 | blood | UM | A/G |
| sample17 | blood | UM | A/G |
| sample18 | blood | UM | G/G |
| sample19 | blood | UM | G/G |
| sample20 | blood | UM | G/G |
| sample21 | blood | UM | G/G |
| sample22 | blood | UM | G/G |
| sample23 | blood | UM | G/G |
| sample24 | blood | UM | G/G |
| sample25 | blood | UM | G/G |
| sample26 | blood | UM | G/G |
| sample27 | blood | UM | G/G |
| sample28 | blood | UM | G/G |
| sample29 | blood | UM | G/G |
| sample30 | blood | UM | G/G |
| sample31 | blood | UM | G/G |
| sample32 | blood | UM | G/G |
| sample33 | blood | UM | G/G |
| sample34 | blood | UM | G/G |
| sample35 | blood | UM | G/G |
| sample36 | blood | UM | G/G |
| sample37 | blood | UM | G/G |
| sample38 | blood | UM | G/G |

Abbreviations: UM/BEB, University of Miami's Brain Endowment Bank; Emory, Emory University Goizueta Alzheimer's Disease Research Center Brain Bank; UM, University of Miami

**Supplementary Table 2.** Linkage disequilibrium analysis showing variants with $r^2$ >0.75 with rs10423769 in ADSP 36k Release and the frequency in 38 samples using LRWGS.

| CHR | Position | rs_id | $r^2$ in ADSP 36k | Frequency in LRWGS (n=38) |
|---|---|---|---|---|
| 19 | 43100141 | rs13344412 | 0.985596 | 0.282 |
| **19** | **43100929** | **rs10423769** | **1** | **0.269** |
| 19 | 43107376 | rs10416442 | 0.994283 | 0.256 |
| 19 | 43109437 | rs10418908 | 0.996912 | 0.269 |
| 19 | 43110194 | rs10426779 | 0.996914 | 0.269 |
| 19 | 43111648 | rs10412855 | 0.997354 | 0.269 |
| 19 | 43111823 | rs10412948 | 0.987293 | 0.269 |
| 19 | 43111824 | rs10413134 | 0.989898 | 0.269 |
| 19 | 43112698 | rs28521996 | 0.994283 | 0.269 |
| 19 | 43113370 | rs10422131 | 0.993395 | 0.269 |
| 19 | 43113989 | rs10402412 | 0.997354 | 0.269 |
| 19 | 43115785 | rs28463422 | 0.954943 | 0.269 |
| 19 | 43115960 | rs28496375 | 0.989493 | 0.269 |
| 19 | 43117612 | rs10425962 | 0.994283 | 0.269 |
| 19 | 43119195 | rs10426472 | 0.797423 | 0.269 |
| 19 | 43122350 | rs11880518 | 0.795646 | 0.231 |
| 19 | 43122490 | rs11880540 | 0.797494 | 0.231 |
| 19 | 43122910 | rs145564654 | 0.787618 | 0.231 |
| 19 | 43123128 | rs138103473 | 0.793468 | 0.231 |
| 19 | 43123613 | rs116423714 | 0.796079 | 0.231 |
| 19 | 43124823 | rs11879815 | 0.783752 | 0.231 |
| 19 | 43128072 | rs114661163 | 0.784624 | 0.218 |
| 19 | 43128687 | rs77244015 | 0.786836 | 0.231 |
| 19 | 43129746 | rs16989239 | 0.784624 | 0.231 |
| 19 | 43132544 | rs116595856 | 0.785961 | 0.231 |
